# Utility of the Recombinase Driver CX3CR1::Cre Rat Strain to Evaluate Microglial Contributions to Nicotine Addiction

**DOI:** 10.1101/2025.03.18.643780

**Authors:** Ashley White, Ashley J. Craig, Daryl Richie, Christa L. Corley, Emma O. Bondy, Percell T. Kendrick, Haley McManus, Safiyah M. Sadek, Michael D. Scofield, Cassandra D. Gipson

## Abstract

Smoking remains a leading preventable cause of death, and nicotine is the primary substance responsible for maintaining use of tobacco products. Preclinical rodent models have shown that neuroimmune signaling is dysregulated by nicotine self-administration (SA) within the nucleus accumbens core (NAcore), which is a key region within the mesolimbic brain reward pathway. Microglia are the resident brain immune cell and prior studies have shown that they play an important role in nicotine-related behaviors. However, while there are transgenic mouse lines that allow for specific evaluations of microglia to neurobiology and behavior, there are fewer tools available for rats as a model species thus limiting our ability to evaluate specific contributions of microglia to nicotine SA. Using transgenic rats expressing Cre under the control of the CX3CR1 promoter bred on a Long Evans (LE) background, we show that NAcore microglia can be specifically transduced with Designer Receptors Exclusively Activated by Designer Drugs (DREADDs). We further show that CX3CR1::Cre rats readily self-administer nicotine, and display a characteristic extinction curve that is not different from outbred LE or Cre-negative littermates. Together, these validation studies lay the foundation for future use of this transgenic rat line to evaluate the specific contributions of microglia in the brain to neurobehavioral underpinnings of nicotine addiction.

## Introduction

The immunomodulatory effects of nicotine have been clinically (Sher et al. 1999) and preclinically (Li et al. 2021; Namba et al. 2019; Shi et al. 2009) documented, however, the impacts of chronic, volitional nicotine use on neuroimmune signaling is a field in its infancy. Microglia are resident brain immune cells which regulate adaptive and innate immune responses (Paolicelli et al. 2022), regulate synaptic plasticity (Aramideh et al. 2021; Bessis et al. 2007; Paolicelli and Ferretti 2017), and are a part of the neurovascular unit (Liu et al. 2020). Recent studies have implicated microglia in nicotine addiction processes as microglia within the reward pathway and specifically the nucleus accumbens (NA) play a large role in regulating nicotine-related neurobehavior (Kumar et al. 2024; Adeluyi et al. 2019; Soares and Picciotto 2024). However, our ability to specifically manipulate microglia to determine causal roles of these brain immune cells in nicotine addiction processes remains limited.

The current neuroscience toolbox for evaluating microglia includes viruses, chemicals to deplete them, and transgenic rodent models. Viruses that have been utilized in prior studies include Designer Receptors Exclusively Activated by Designer Drugs (DREADDs) which incorporate adeno-associated viral vectors (AAVs) with promoters that are thought to be microglia-specific (e.g., CD68; (Grace et al. 2016), and AAV variants that allow *in vitro* and *in vivo* microglial transduction through directed evolution of the AAV capsid protein to increase efficiency of genetic payload into microglia (Lin et al. 2022). Notably, the pAAV-CD68-hM4D-mCherry (Gi) and pAAV-CD68-EGFP DREADD utilized in (Grace et al. 2016) are commercially available but untested in their ability to penetrate microglia within the brain as that study infused DREADDs intrathecally. The AAV variants which were discovered a few years later to infect microglia were also found to infect neurons (Lin et al. 2022), thus making it difficult to tease apart microglial versus neuronal contributions to neurobiology and behavior.

As mentioned above, other methodologies to evaluate contributions of microglia to brain function include depletion methodologies with either a Cx3cr1-Dtr rat, which allows for short-term microglia depletion (De Luca et al. 2019), or with colony-stimulating factor–1 receptor (CSF1R) inhibitors (e.g., PLX5622 or PLX3397), which deplete a proportion of microglia as well as other affect peripheral cells (Green et al. 2020) when administered in chow. An advantage of this technique is that microglia can repopulate (Wickel et al. 2024), allowing for bidirectional evaluations of microglia to neurobiology and behavioral outcomes. However, microglia depletion is non-specific and can have off target effects, limiting the cell-type specificity of this approach. An additional methodology includes frequently utilized transgenic mouse strains which have been used primarily in other fields of research. Transgenic mice, including the recombinase driver CX3CR1-Cre strains (Zhao et al. 2019), have been used to visualize or manipulate microglia in the context of other pathologies (Wieghofer and Prinz 2016). Endogenous CX3CR1 is a microglia-specific marker, and it is a receptor with the endogenous ligand fractalkine (Nishiyori et al. 1998). However, there has been some debate regarding specificity of CX3CR1 to microglia, as there are reports that CX3CR1 expression occurs in other cell types depending on the tissue type (Lee et al. 2018), or co-localization of CX3CR1 with glial fibrillary acidic protein (GFAP; (Peng et al. 2017), a marker of astrocyte function (Middeldorp and Hol 2011), although this has not been evaluated within the brain reward pathway to our knowledge. Some studies have also reported that CX3CR1::cre in mice is not specific to microglia, with significant leakage into neurons (Haimon et al. 2018; Hwang et al. 2017; Zhang et al. 2018). While there are two constitutive and two tamoxifen-inducible CX3CR1::cre mouse lines (Gong et al. 2007; Parkhurst et al. 2013; Yona et al. 2013), there is only one CX3CR1::cre rat line which has never been utilized to evaluate microglia contributions to addiction neuroscience and has never been demonstrated to allow for microglia-specific infection.

A large majority of studies modeling nicotine addiction utilize self-administration (SA), which allows for evaluations of volitional nicotine use and is most frequently implemented in rat models (Corley et al. 2024). Studies also frequently incorporate extinction training following SA to evaluate the ability to learn to withhold the prepotent response to receive the drug and is typically conducted under withdrawal conditions (Namba, et al., 2019). While some studies incorporate mouse SA to model aspects of nicotine addiction (Fowler et al. 2011; Fowler and Kenny 2011), use of mice in nicotine SA is limited to a small number of researchers and can present challenges with catheter patency among other issues. While all currently available methodologies to evaluate microglia have limitations with some noted above, the microglia addiction field has a significant barrier in that there are no validated rat strains for use in SA procedures. Here, we utilized a commercially available CX3CR1::cre transgenic rat strain (LE-Tg(OTTC1005) CX3CR1-Cre-ERT2), which is a tamoxifen-inducible Cre recombinase strain based on a Long Evans background. The goal of the current study was therefore to determine if this rat strain (1) displays microglia-specific expression of viral vectors within the NAcore without infecting surrounding neurons, and (2) can acquire nicotine SA and extinction (3) similar to CX3CR1::cre (-) littermates and wildtype Long Evans rats.

## Methods

### Subjects

Breeder male LE-Tg(OTTC1005) CX3CR1-Cre-ERT2 rats were purchased from the Rat Resource and Research Center (RRRC) and bred in-house with Long Evans females purchased from Charles River. Rats were housed in a temperature and humidity-controlled animal facility within the Division of Laboratory Animal Resources (DLAR) at the University of Kentucky. The colony room was on a 12:12 hr light:dark reverse light cycle and rats in the breeding colony had *ad libitum* access to food and water. Rats in nicotine SA were either CX3CR1::Cre (+; N=50), CX3CR1::Cre (-; N=15), or purchased from Charles River as an outbred strain control (N=37). Male and female rats were 60-90 days old at the beginning of the study and were food restricted to 85% of free feed weight throughout SA experimentation. All animal use practices were approved by the Institutional Animal Care and Use Committee (IACUC) of the University of Kentucky (UK; Protocol #2020-3438). 19 CX3CR1::Cre (+), 3 CX3CR1::Cre (-), and 2 Long Evans rats were excluded from the study due to catheter patency issues.

### CX3CR1::Cre Genotyping

Ear punches were collected from all animals on post-natal day 21. In separate 1.5 mL microcentrifuge tubes, ear punch tissues were incubated in lysis buffer and Proteinase K (Invitrogen, ∼2 mg/ml, 24 µl total volume) at 55°C for 30 minutes, vortexed, and pelleted (18,000 x g). Samples were incubated for an additional 30 min at 55°C, vortexed, then resuspended in distilled water up to 200 µl total volume, and pelleted (18,000 x g). The samples were heated at 100°C for 10 minutes, cooled to room temperature, and pelleted at 18,000 x g for 20 minutes at 4°C. Next, 100 µl of the supernatant containing the genomic DNA was transferred to a new 1.5 ml microcentrifuge tube, and standard PCR amplification was used for genotyping.

Subjects were genotyped using the following primers: CX3CR1 F103289 (5’-TCAGGGTGG CCCATAACCAC-3*’*) and CreETT2 R713 (5’-AGGTAGTTATTCGGATCATCAGCTTAC AC-3’) which produce a 1017 bp amplicon spanning the 5’ end of the CX3CR1-Cre-ERT2. Primers CX3CR1 F103289 (5’-TCAGGGTGGCCCATAACCAC-3*’*) and CREERT2 R246 (5’-CAGTGA AGCTTCGATGATGGG C-3’), which produce a 550 bp amplicon spanning the 5’ end of the CX3CR1-Cre-ERT2. Primers CreERT2 F1206 (5’-CAGTGAAGCTTCGATGATGGGC-3’) and CX3CR1 R104157 – 5’ – GCCAGATTTCCTGCAGGACCT-3’), which produce a 1553 bp amplicon spanning the 3’ end of the CX3CR1-Cre-ERT2. All animals used were confirmed CX3CR1::cre transgene-positive.

### Surgical Procedures

Rats included in the immunohistochemistry study underwent surgery for viral infusions targeting the NAcore as previously described (Leyrer-Jackson et al. 2021). Rats included in the nicotine SA study underwent jugular catheter surgery and virus infusion surgery. All rats were anesthetized using ketamine hydrochloride (80–100□mg/kg, i.m.) and xylazine (8□mg/kg, i.m.) and underwent surgical implantation of intravenous jugular catheters as well as stereotaxically infused viral delivery within the NAcore. Intravenous jugular catheters (made from polyurethane tubing; BTPU-040; Instech) were inserted 2.5–3□cm into the right jugular vein and were threaded subcutaneously to the posterior side of the animal where it was connected to an indwelling back port (Instech). Dental cement (SNAP or Ortho-Jet) was used to adhere the catheter to the port. The indwelling port was sutured using 4–0 vicryl braided suture (Ethicons) subcutaneously □2□cm caudal from the shoulder blades. Following jugular vein catheterization, animals were immediately transferred to a rat stereotaxic frame and NAcore guide cannulae were bilaterally aimed at the NAcore according to a stereotaxic atlas (Paxinos and Watson 2007). Viral vectors encoding DREADDs for microglia infection were infused into the NAcore of CX3Cr1::cre animals: pAAV-EF1a-DIO-hM4D(Gi)-mCherry (inhibitory; titer: 1.2□×□1013 vg/ml; Addgene, #50461), pAAV-EF1a-DIO-hM3D(Gq)-mCherry (excitatory; titer: 1.3□×□1013 vg/ml; Addgene, #50460; packaged into an AAV5 vector by UNC Vector Core), or pAAV5-EF1a-DIO-mCherry (control; i.e., only mCherry-expressing; titer: 1.5□×□1013 vg/ml; Addgene, #44362) at a volume of 0.5□μl per hemisphere. Ef1a was selected as the promoter because it is abundantly expressed in all cell types (Wakabayashi-Ito and Nagata 1994), it is double floxed, and both Gq and Gi coupled-versions are commercially available. Rats were immediately administered cefazolin (100□mg/kg, i.v.) and heparin (10 U/ml, i.v.) and for seven consecutive days during the recovery period. Meloxicam (1□mg/kg, i.v.) was given immediately and for the first 3□d of the recovery period. For rats in the SA study, heparin (10 U/ml, i.v.) was administered daily. All rats received 4-hydroxytamoxifen (20 mg/kg, I.P.; (Hughes et al. 2024) 3 weeks following virus infusion and 2 days prior to experimentation.

### Food Training Procedures

All rats in the nicotine SA study (N = 30) underwent food training on the sixth day of postoperative care. Food restriction to maintain 85% of free feed weight was implemented a minimum of 2 h before food training initiation and was maintained for the duration of the experiment. Food training sessions were 15 h duration, and one active lever press resulted in the delivery of one food pellet [fixed-ratio-1 (FR1), schedule of reinforcement; Bio-Serv, 45□mg/pellet) Concurrently, one food pellet was delivered every 20 min regardless of response. Light and tone stimuli were not paired with pellet administration. Food training criteria was set to 2:1 active to inactive lever presses throughout the session and a minimum of 200 active lever presses. Rats that did not pass the initial food training session completed a second session before beginning SA procedures (N=10). Rats unable to pass food training procedures were excluded from the study (N = 8). All food training, SA, and extinction sessions were conducted in modular operant conditioning chambers (13 ENV-008, 15 ENV-007; Med Associates), which have been previously described in detail (Sadek et al. 2024; Maher et al. 2023; Leyrer-Jackson et al. 2021).

### Nicotine SA Procedures

Nicotine infusions (0.06□mg/kg/infusion) were paired with a compound stimulus (light+tone) and was followed by a 20-s timeout period. Nicotine was delivered across a 5.9-s duration, at a total volume of 0.1 ml of nicotine was administered per infusion. Infusions were paired with cue lights, located above each lever, and a 2900-Hz tone was presented throughout the duration of the nicotine infusion. Active lever presses during the timeout period were recorded, but did not result in additional nicotine infusions. An inactive lever was extended at all times and responses were recorded; however, presses yielded no programmed consequences or rewards. Session duration was 2 h in length and a FR1 schedule of reinforcement was used. All animals were required to complete a minimum of 10 sessions before moving into extinction with the following criteria: ≥10 nicotine infusions obtained and ≥2:1 active/inactive lever press ratio. Additionally, extinction is defined either by (1) the lack of delivery of a reinforcer that was previously delivered after a response (e.g., responses on the active lever no longer result in nicotine infusions), or (2) as the absence of a contingency between response and reinforcer (e.g., the nicotine infusion occurs regardless of an animals’ response; see Rescorla and Skucy, 1969; Lattal and Lattal, 2012). Given that the reinforcer here (the nicotine infusion) is not delivered following a time-out active lever press, the latter definition applies to these types of responses and are therefore distinct from active lever presses that directly result in the initiation of a nicotine infusion. Thus, these two types of lever presses were analyzed separately in the current study.

Following 10 non-consecutive criteria-making sessions, rats were moved to the extinction phase which took place in the same boxes as self-administration. During the extinction phase, active lever presses no longer resulted in nicotine infusions or associated cues. A minimum of 14 2-h extinction sessions was required for each animal. Extinction criteria were set at <30 active lever presses on the last day of extinction. Eight animals were removed from the current study because of not meeting self-administration criteria (*N*□=□5) or lack of DREADD expression/improper cannula placement (*N*□=□3).

### Immunohistochemistry

Rats were anesthetized with an overdose of euthasol (excess of 0.25 ml/kg i.p.) and transcardially perfused with 0.01 M PBS followed by 4% paraformaldehyde. Brains were then coronally sliced on a vibratome at 60 µm for immunohistochemistry staining. All antibodies and dilutions are listed in **Table 1**. To determine if the virus infected microglia cells in the NAcore, NAcore sections were first incubated in anti-Iba-1 primary antibody that was diluted in 1% BSA, 0.3% Triton, .05% azide, and 0.01 M PBS for 24 hours at room temperature on a shaker. Sections were then rinsed with 0.01 M PBS three times for 30 minutes each then incubated in a secondary antibody conjugated to Alexa Fluor 488 for 2 hours. Sections were mounted onto glass slides and coverslipped using an anti-fade mounting medium. Next, to determine if the virus infected microglia that expressed the cre (+) genotype, NAcore sections were simultaneously stained with 2 primary antibodies (anti-Iba1, and anti-CX3CR1) that were diluted in 1% BSA, 0.3% Triton, .05% azide, and 0.01 M PBS for 24 hours at room temperature on a shaker. Sections were then rinsed and incubated in secondary antibodies conjugated to Alexa Fluor 488 and Alexa Fluor 594 for 2 hours. Sections were mounted onto glass slides and coverslipped using an anti-fade mounting medium containing DAPI to visualize cell bodies. Next to show colocalization of infected microglia without infected neurons, NACore sections were incubated in 2 primary antibodies (anti-Iba1, and anti-NeuN) that were diluted in 1% BSA, 0.3% Triton, .05% azide, and 0.01 M PBS for 24 hours at room temperature on a shaker. Sections were then rinsed and incubated in secondary antibodies conjugated to Alexa Fluor 488 and Alexa Fluor 594 for 2 hours. All confocal micrographs were captured using a Nikon A1R Inverted Confocal Microscope.

**Table 1.**
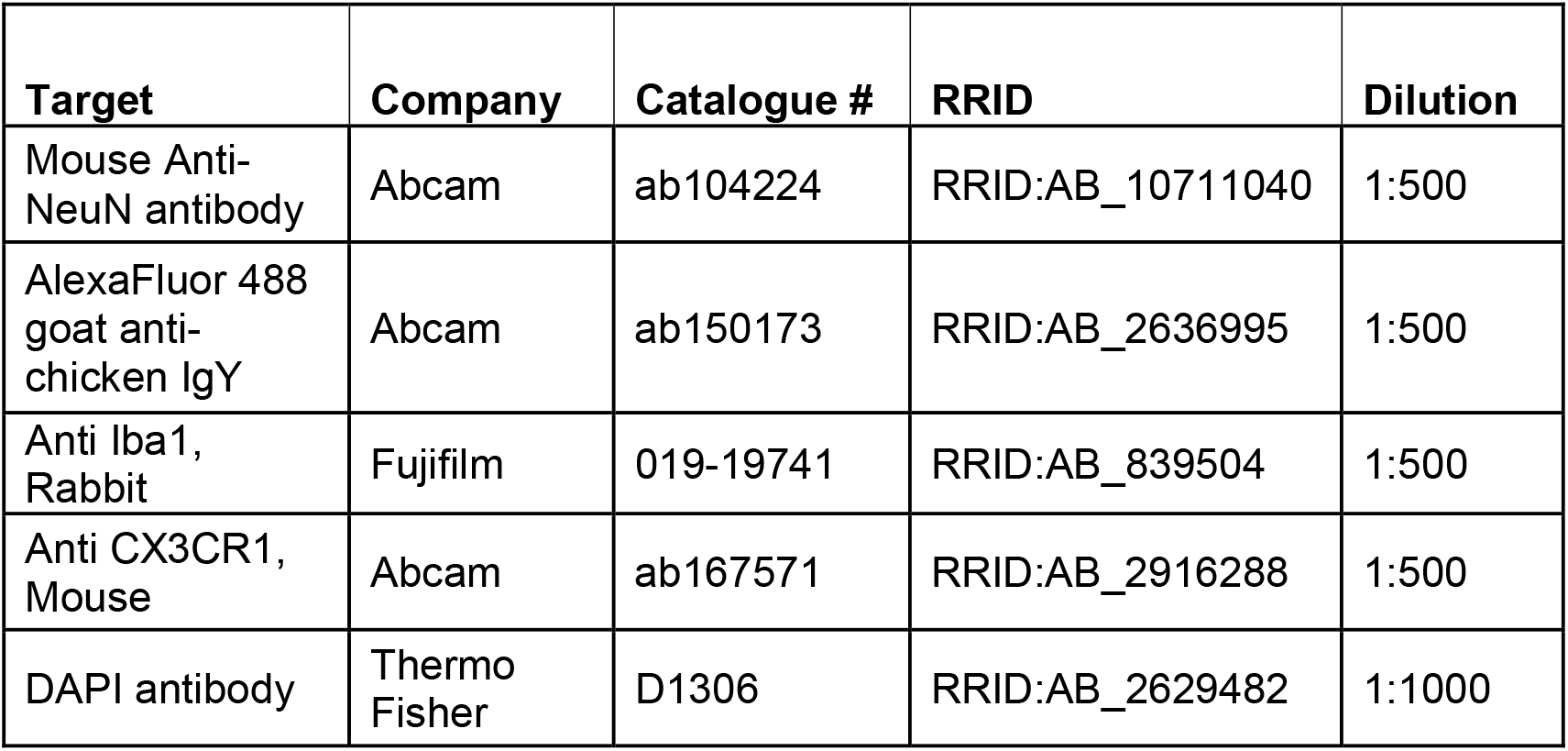
Antibodies for Immunohistochemistry.

### Statistical Analysis

Food training lever press data, drug infusions, and lever press discrimination (active, inactive) across sessions were analyzed using linear mixed effects (LME) modeling (JMP software from SAS). Tukey’s HSD was used as post-hoc pairwise comparisons on nominal variables where appropriate (α = 0.05). Subject was treated as a random factor and session was treated as a continuous factor whereas group was evaluated as a nominal variable in all relevant analyses. Model fits were conducted to determine the best model fit for the datasets where appropriate, and the models with the lowest Akaike information criterion (AICc) were selected. Lever discrimination was assessed as the ratio of active/active+inactive lever presses (including infusion-earning and timeout active lever presses, as well as all inactive lever presses).

## Results

### Cre-Dependent Viruses Express Within NAcore Microglia

Immunohistochemistry was first conducted to determine if the fluorescent mCherry tag from the viruses expressed in NAcore microglia following intracranial viral delivery. Images were taken using a Nikon A1R Confocal Microscope. **Figure 1A** shows that mCherry from the Gi viral construct was successfully observed within NAcore Iba1-(+) cells. Similarly, we found co-localization of mCherry with NAcore Iba1-(+) cells with the Gq construct (**Figure 1B**). To determine specificity to CX3CR1-expressing cells, we next evaluated co-localization of mCherry from the Gi construct with CX3CR1 and Iba-1. Notably, cells that were mCherry-(+) expressed both microglia markers (**Figure 1C**), demonstrating that the CX3CR1::cre transgenic rat strain allowed for AAV infection of microglia. Finally, we confirmed somatic expression of mCherry with co-localization of the fluorescent tag with DAPI, which visualizes nuclear DNA (Tarnowski et al. 1991); **Figure 1D**). Next, we determined if mCherry from the Gi construct co-localized with the neuronal marker NeuN. As shown in **Figure 1E**, there was no co-localization of mCherry with NeuN, demonstrating specificity of mCherry to microglia within the NAcore.

**Figure 1.**
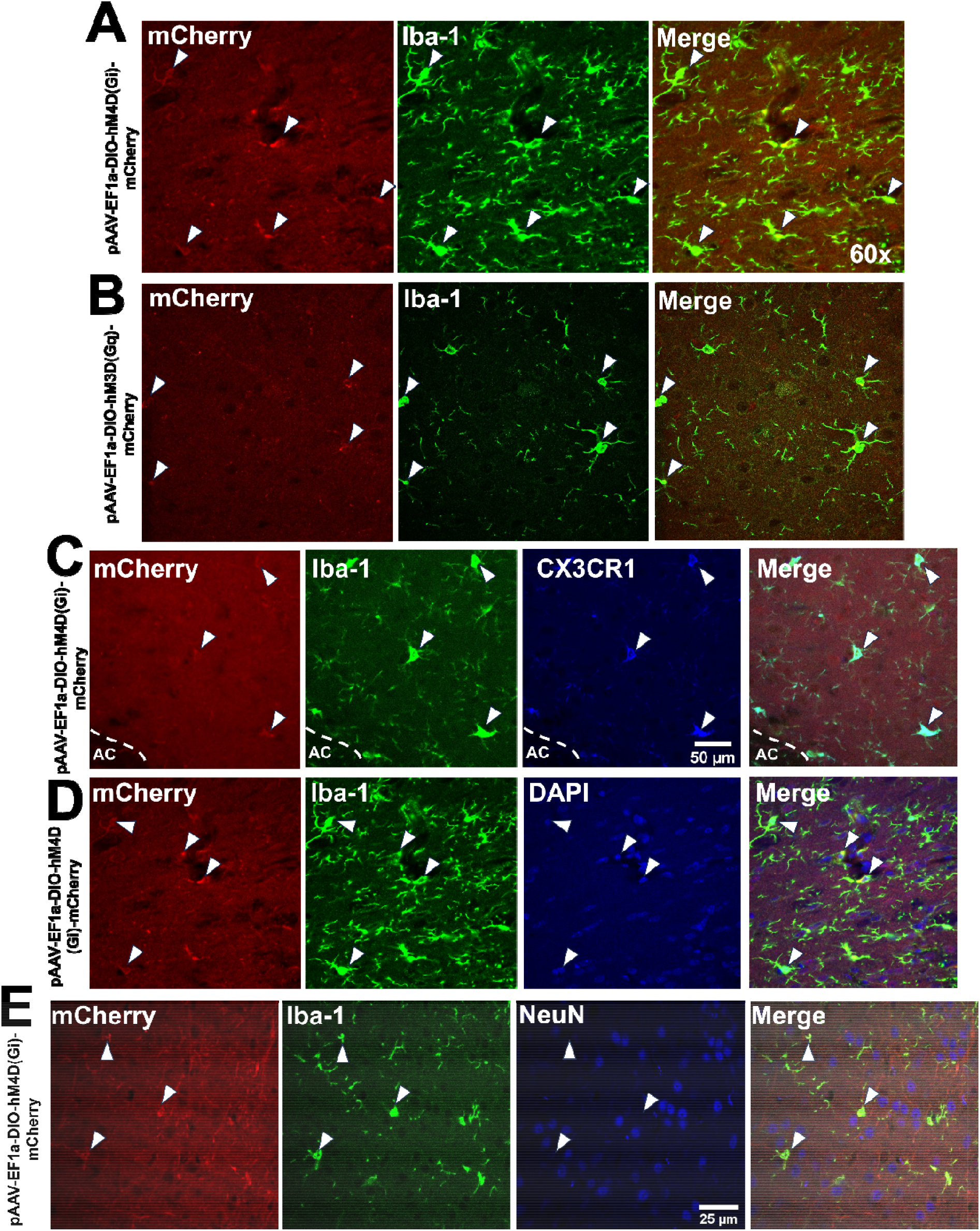
NAcore Microglia but not Neurons Are Transduced with AAV Constructs. Representative immunohistochemical images from animals receiving viral infusions are shown. Slices were stained with either whole cell microglial marker, Iba1; Cre-recombinase marker, mCherry; microglial receptor, CX3CR1; cell soma marker, DAPI, and/or neuronal cell marker, NeuN. Merged images show overlap of microglial and viral markers, indicating infection of microglia with viral constructs without neuronal infection. AC = anterior commissure, denoted by a dotted line.

### CX3CR1::cre Rats Acquire Nicotine SA and Extinction

First, baseline nicotine consumption was compared during nicotine SA acquisition as a function of virus groups following viral administration (see **Figure 2A** for an experimental timeline). The best model fit excluded sex in the model (ΔAICc = 31.9, relative to the next best model fit, which included both sex and group as a variable). LME revealed no significant main effects group (F_2,2_ = 0.6081, *p*>0.05), session (F_1,21_ = 1.0336, *p*>0.05), or no group x session interactions (F_2,21_ = 0.4445, *p*>0.05) (**Figure 3B**). These results indicate that virus administration did not change nicotine SA acquisition rates regardless of the construct type. When nicotine consumption was summed across experimentation, there were no group differences in the total amount of nicotine consumption across the 15-day SA experiment (F_2,21_ = 1.2943, *p*>0.05).

**Figure 2.**
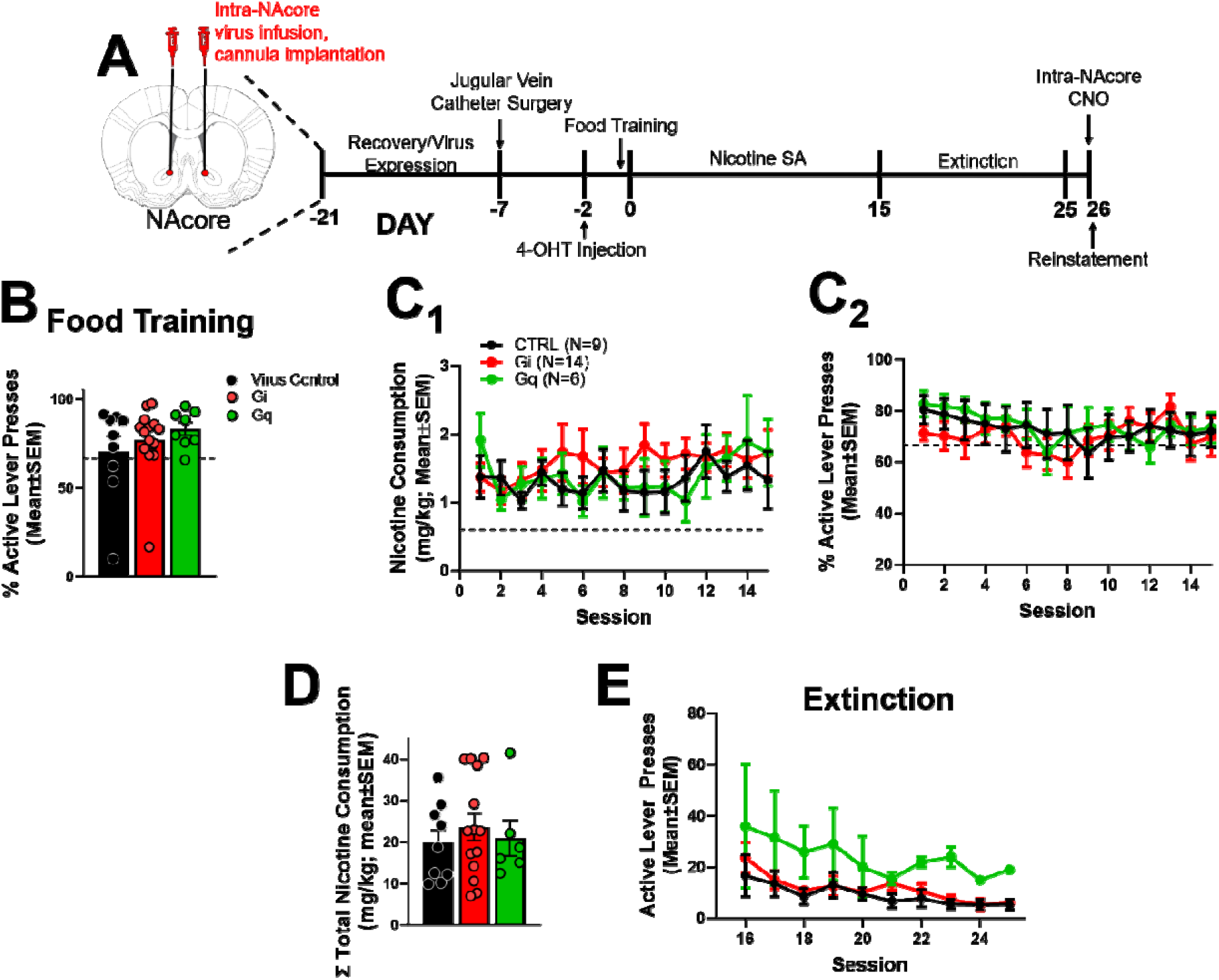
CX3CR1::cre Rats Acquire Nicotine SA and Extinction. (A) Experimental timeline for CX3CR1::cre rats. (B) Food training where animals met a 2:1 active to inactive lever pressing ratio compared across virus groups with no significant differences *(p>0*.*05)*. (C_1-2_) CX3CR1::cre rats acquired lever responding to the active lever to receive IV infusions of nicotine (0.06mg/kg/infusion) with no differences in lever discrimination. (D) CX3CR1::cre rats had no significant differences in total nicotine consumption across the 15 day timeline. (E) CX3CR1::cre rats showed a decline in lever pressing behavior during extinction training to < 30 lever presses per session.

**Figure 3.**
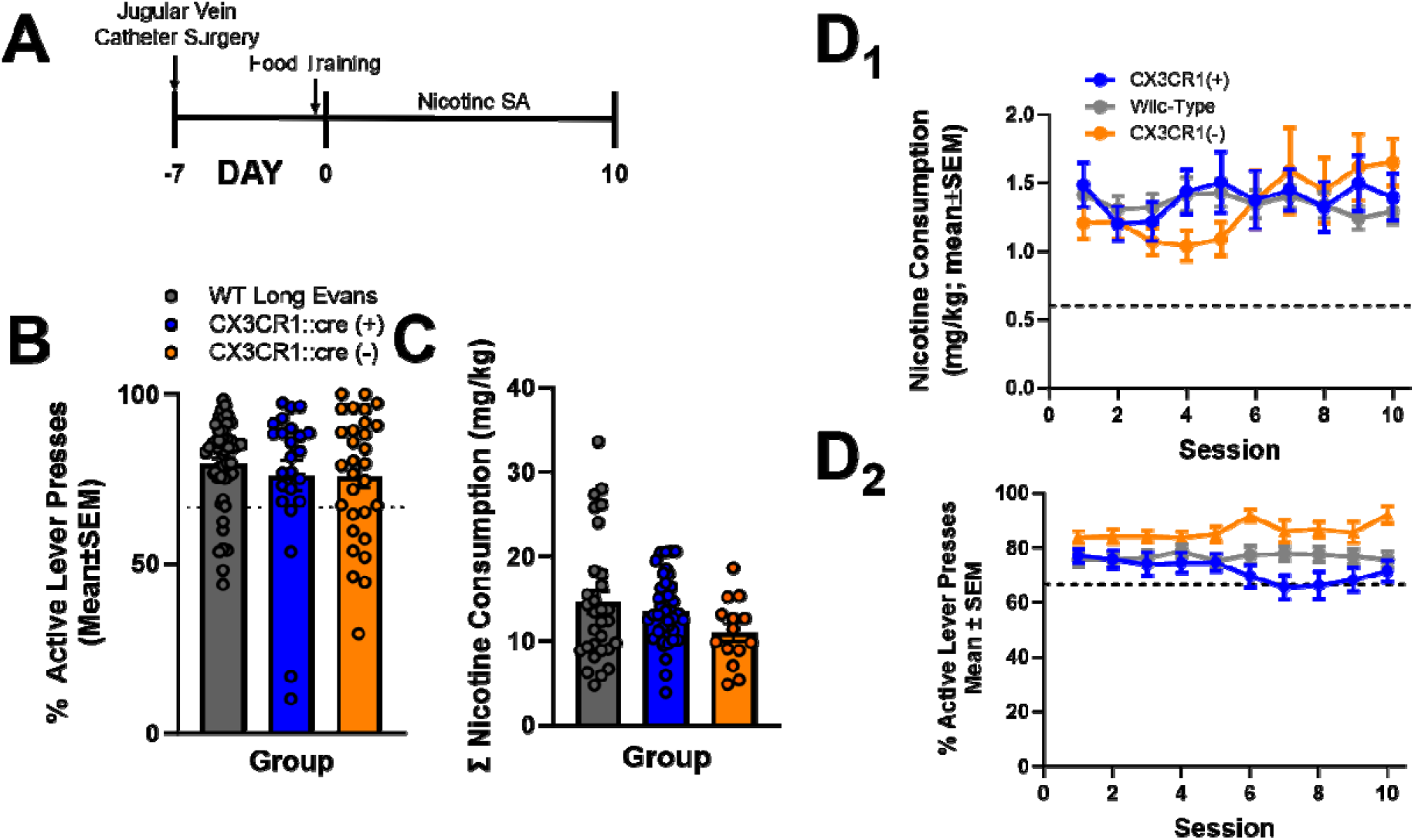
CX3CR1::cre Rats Acquire Nicotine SA similarly to Cre-negative Littermates and Wild-Type rats. (A) Experimental timeline for cre-negative littermates and ordered wild-type rats from Charles River. (B) Food training where animals met a 2:1 active to inactive lever pressing ratio compared across strain groups with no significant differences *(p>0*.*05)*. (C) Rats had no significant differences in total nicotine consumption across the 10 day timeline across strains. (D_1-2_) CX3CR1::cre rats acquired lever responding to the active lever to receive IV infusions of nicotine (0.06mg/kg/infusion) compared to cre-negative littermates and wild-type ordered animals with a significant difference in lever discrimination between strains (p<0.05).

Next, data were collapsed across virus type and compared to CX3CR1::cre (-) littermates, as well as purchased Long Evans rats. The best model fit excluded sex in the model (ΔAICc = 44.2, relative to the next best model fit, which included both sex and group as a variable). First LME was conducted on lever discrimination during food training as a function of strain. No main effect of strain (F_1,68_ = 0.44, *p*>0.05 was found, indicating that all rats acquired food training similarly with no strain differences in this effect. Baseline nicotine consumption was compared during nicotine SA acquisition as a function of strain groups (see **Figure 3A** for an experimental timeline). LME revealed no significant main effects of group (F_2, 64_ = 1.1342, *p*>0.05), or session (F_1, 64_ = 3.0377, *p*>0.05) or group x session interactions (F_2, 64_ = 2.2673, *p*>0.05) (**Figure 3B**). These results indicate that the strain did not change nicotine SA acquisition rates regardless of the genetic strain used. LME was conducted on lever discrimination during nicotine acquisition as a function of strain. A significant main effect of strain (F_2,64.02_ = 5.3284, *p<*0.05) was found, however, all groups met the 2:1 active to inactive lever pressing ratio. When nicotine consumption was summed across experimentation, there were no group (F_2,77_ = 1.7039, *p>*0.05) differences in the total amount of nicotine consumption across the 10-day SA experiment.

## Discussion

Here we show that the CX3CR1::cre transgenic rat strain can be used to specifically transduce DREADDs in microglia within a key region of the brain reward pathway, the NAcore, and these rats can acquire intravenous nicotine SA and also undergo successful nicotine extinction training. We compared acquisition curves across virus type and across strain with no differences in either analyses. As noted previously, evaluations of microglia in addiction neurobiology have been limited due to the tools available to study this cell type in rat models. Our current data reported here support the use of this transgenic rat strain in the addiction neuroscience field to evaluate microglia contributions to substance use, and particularly, in the nicotine addiction field.

While the current study demonstrates the utility of this rat strain for studying aspects of addiction neuroscience, there are notable limitations that must be outlined. It is important to note that the current study did not functionally evaluate the impacts of chemogenetic activation or inhibition on NAcore microglia, including evaluations of transcriptomic, epigenetic, metabolomic, or proteomic changes induced by activation of the DREADD receptors. This is important because the microglia field has recently shifted from a dichotomous thinking of “activated” versus “resting” microglial states to a more complex multidimensional approach to microglial states (Paolicelli et al. 2022). It is also unknown how muscarinic DREADD receptors expressed on CX3CR1(+) microglia may shift these dynamic aspects of these immune cells. One study did find that activation of Gi DREADDs expressed in microglia of CX3CR1^creER/+^:R26^LSL- hM4Di/+^ transgenic mice inhibited chronic pain and spinal nerve transection-induced microglial reactivity (Yi et al. 2021). This study also found that the Gi DREADD reduced interferon regulatory factor 8 (IRF8) and interleukin (IL)-1β expression in Iba1(+) cells, although this study did not confirm that all microglia were DREADD(+). Further, while this rat strain allows for infection of CX3CR1(+) microglia, this methodology requires these cells to be virally infected with the construct. Here we did not find that all microglia within the NAcore were mCherry(+), indicating that some microglia within the area of viral delivery were not infected with the constructs. This could either be due to (1) too little infused volume of the virus to perfuse through the visualized area of the NAcore, (2) some microglia did not express cre, or (3) some microglia were able to mount an immune response from the viral constructs. It is also important to note that this rat strain requires tamoxifen injections. Given that tamoxifen is a selective estrogen receptor modulator and may have sex-specific antiestrogenic effects (Rochefort and Borgna 1981) that could interfere with experimental outcomes, future studies incorporating CX3CR1::cre rats should take this into account in experimental timelines. Here, rats were administered tamoxifen several weeks prior to quantification of microglia morphology, and during the early part of nicotine SA acquisition. Importantly, tamoxifen has a half-life of 10.3 hrs in rats (Robinson et al. 1991), and five elimination half-lives are required for approximately 97% of the bioavailable dose of a drug to be eliminated from the body (here, 51.5 hrs would be required; (Nnane 2005). Given this and that no sex differences were found in the nicotine SA acquisition curves, it is unlikely that tamoxifen impacted behavioral outcomes here. As well, another study administered 4-OHT immediately prior to cocaine conditioning and observed measurable cocaine place preference in mice (Hughes et al. 2024). Finally, it is not clear if CX3CR1::cre rat line has altered constitutive CX3CL1/CX3CR1 signaling, which is important in neuron/microglia crosstalk in brain injury and inflammation (Pawelec et al. 2020). Despite these caveats, this rat strain may allow for more granular *in vivo* evaluations of microglia to glutamate dyshomeostasis induced by addictive drugs (Scofield et al. 2016) including nicotine (Gipson et al. 2013), as well as cell type-specific dissections of glial contributions to the neurobehavior of nicotine addiction.

## Conclusions

Here, we have shown that the inducible recombinase driver CX3CR1::Cre rat strain can be successfully used in a model of nicotine SA. CX3CR1::Cre rats readily self-administer nicotine, while the viral constructs successfully infected microglia without neuronal co-infection within the NAcore, eliminating one of the main issues with other methodologies utilizing viral constructs. While mouse nicotine SA can be conducted successfully (Souter, et al., 2022), the majority of SA studies utilize rats as the model rodent species, thus this rat will provide researchers with the toolbox to readily manipulate microglia in rats undergoing SA of addictive drugs. Understanding these contributions will provide insights on the neuroimmune landscape of nicotine addiction and how immune modulation in different areas of the brain affect behavior.

## REFERENCES

Adeluyi A, Guerin L, Fisher ML, Galloway A, Cole RD, Chan SSL, Wyatt MD, Davis SW, Freeman LR, Ortinski PI, Turner JR (2019) Microglia morphology and proinflammatory signaling in the nucleus accumbens during nicotine withdrawal. Sci Adv 5 (10):eaax7031. doi:10.1126/sciadv.aax7031

Aramideh JA, Vidal-Itriago A, Morsch M, Graeber MB (2021) Cytokine Signalling at the Microglial Penta-Partite Synapse. Int J Mol Sci 22 (24). doi:10.3390/ijms222413186

Bessis A, Bechade C, Bernard D, Roumier A (2007) Microglial control of neuronal death and synaptic properties. Glia 55 (3):233–238. doi:10.1002/glia.20459

Corley C, Craig A, Sadek S, Marusich JA, Chehimi SN, White AM, Holdiness LJ, Reiner BC, Gipson CD (2024) Enhancing translation: A need to leverage complex preclinical models of addictive drugs to accelerate substance use treatment options. Pharmacol Biochem Behav 243:173836. doi:10.1016/j.pbb.2024.173836

De Luca SN, Sominsky L, Soch A, Wang H, Ziko I, Rank MM, Spencer SJ (2019) Conditional microglial depletion in rats leads to reversible anorexia and weight loss by disrupting gustatory circuitry. Brain Behav Immun 77:77–91. doi:10.1016/j.bbi.2018.12.008

Fowler CD, Kenny PJ (2011) Intravenous nicotine self-administration and cue-induced reinstatement in mice: Effects of nicotine dose, rate of drug infusion and prior instrumental training. Neuropharmacology 61 (4):687–698. doi:10.1016/j.neuropharm.2011.05.012

Fowler CD, Lu Q, Johnson PM, Marks MJ, Kenny PJ (2011) Habenular alpha5 nicotinic receptor subunit signalling controls nicotine intake. Nature 471 (7340):597–601. doi:10.1038/nature09797

Gipson CD, Reissner KJ, Kupchik YM, Smith AC, Stankeviciute N, Hensley-Simon ME, Kalivas PW (2013) Reinstatement of nicotine seeking is mediated by glutamatergic plasticity. Proc Natl Acad Sci U S A 110 (22):9124–9129. doi:10.1073/pnas.1220591110

Gong S, Doughty M, Harbaugh CR, Cummins A, Hatten ME, Heintz N, Gerfen CR (2007) Targeting Cre recombinase to specific neuron populations with bacterial artificial chromosome constructs. J Neurosci 27 (37):9817–9823. doi:10.1523/jneurosci.2707-07.2007

Grace PM, Strand KA, Galer EL, Urban DJ, Wang X, Baratta MV, Fabisiak TJ, Anderson ND, Cheng K, Greene LI, Berkelhammer D, Zhang Y, Ellis AL, Yin HH, Campeau S, Rice KC, Roth BL, Maier SF, Watkins LR (2016) Morphine paradoxically prolongs neuropathic pain in rats by amplifying spinal NLRP3 inflammasome activation. Proc Natl Acad Sci U S A 113 (24):E3441–3450. doi:10.1073/pnas.1602070113

Green KN, Crapser JD, Hohsfield LA (2020) To Kill a Microglia: A Case for CSF1R Inhibitors. Trends Immunol 41 (9):771–784. doi:10.1016/j.it.2020.07.001

Haimon Z, Volaski A, Orthgiess J, Boura-Halfon S, Varol D, Shemer A, Yona S, Zuckerman B, David E, Chappell-Maor L, Bechmann I, Gericke M, Ulitsky I, Jung S (2018) Re-evaluating microglia expression profiles using RiboTag and cell isolation strategies. Nat Immunol 19 (6):636–644. doi:10.1038/s41590-018-0110-6

Hughes BW, Huebschman JL, Tsvetkov E, Siemsen BM, Snyder KK, Akiki RM, Wood DJ, Penrod RD, Scofield MD, Berto S, Taniguchi M, Cowan CW (2024) NPAS4 supports cocaine-conditioned cues in rodents by controlling the cell type-specific activation balance in the nucleus accumbens. Nature communications 15 (1):5971. doi:10.1038/s41467-024-50099-1

Hwang HW, Saito Y, Park CY, Blachère NE, Tajima Y, Fak JJ, Zucker-Scharff I, Darnell RB (2017) cTag-PAPERCLIP Reveals Alternative Polyadenylation Promotes Cell-Type Specific Protein Diversity and Shifts Araf Isoforms with Microglia Activation. Neuron 95 (6):1334–1349.e1335. doi:10.1016/j.neuron.2017.08.024

Kumar M, Keady J, Aryal SP, Hessing M, Richards CI, Turner JR (2024) The Role of Microglia in Sex-and Region-Specific Blood-Brain Barrier Integrity During Nicotine Withdrawal. Biol Psychiatry Glob Open Sci 4 (1):182–193. doi:10.1016/j.bpsgos.2023.08.019

Lee M, Lee Y, Song J, Lee J, Chang SY (2018) Tissue-specific Role of CX(3)CR1 Expressing Immune Cells and Their Relationships with Human Disease. Immune Netw 18 (1):e5. doi:10.4110/in.2018.18.e5

Leyrer-Jackson JM, Holter M, Overby PF, Newbern JM, Scofield MD, Olive MF, Gipson CD (2021) Accumbens Cholinergic Interneurons Mediate Cue-Induced Nicotine Seeking and Associated Glutamatergic Plasticity. eNeuro 8 (1). doi:10.1523/ENEURO.0276-20.2020

Li H, Peng Y, Lin C, Zhang X, Zhang T, Wang Y, Li Y, Wu S, Wang H, Hutchinson MR, Watkins LR, Wang X (2021) Nicotine and its metabolite cotinine target MD2 and inhibit TLR4 signaling. Innovation (Camb) 2 (2):100111. doi:10.1016/j.xinn.2021.100111

Lin R, Zhou Y, Yan T, Wang R, Li H, Wu Z, Zhang X, Zhou X, Zhao F, Zhang L, Li Y, Luo M (2022) Directed evolution of adeno-associated virus for efficient gene delivery to microglia. Nat Methods 19 (8):976–985. doi:10.1038/s41592-022-01547-7

Liu LR, Liu JC, Bao JS, Bai QQ, Wang GQ (2020) Interaction of Microglia and Astrocytes in the Neurovascular Unit. Front Immunol 11:1024. doi:10.3389/fimmu.2020.01024

Maher EE, White AM, Craig A, Khatri S, Kendrick PT, Jr., Matocha ME, Bondy EO, Pallem N, Breakfield G, Botkins M, Sweatt O, Griffin WC, Kaplan B, Weafer JJ, Beckmann JS, Gipson CD (2023) Synthetic contraceptive hormones occlude the ability of nicotine to reduce ethanol consumption in ovary-intact female rats. Drug Alcohol Depend 252:110983. doi:10.1016/j.drugalcdep.2023.110983

Middeldorp J, Hol EM (2011) GFAP in health and disease. Prog Neurobiol 93 (3):421–443. doi:10.1016/j.pneurobio.2011.01.005

Namba MD, Kupchik YM, Spencer SM, Garcia-Keller C, Goenaga JG, Powell GL, Vicino IA, Hogue IB, Gipson CD (2019) Accumbens neuroimmune signaling and dysregulation of astrocytic glutamate transport underlie conditioned nicotine-seeking behavior. Addict Biol:e12797. doi:10.1111/adb.12797

Nishiyori A, Minami M, Ohtani Y, Takami S, Yamamoto J, Kawaguchi N, Kume T, Akaike A, Satoh M (1998) Localization of fractalkine and CX3CR1 mRNAs in rat brain: does fractalkine play a role in signaling from neuron to microglia? FEBS Lett 429 (2):167–172. doi:10.1016/s0014-5793(98)00583-3

Nnane IP (2005) PHARMACOKINETICS: Absorption, Distribution, and Elimination. In: Paul Worsfold AT, Colin Poole (ed) Encyclopedia of Analytical Science. 2nd edn. Elsevier, pp 126–133. doi:10.1016/B0-12-369397-7/00458-1.

Paolicelli RC, Ferretti MT (2017) Function and Dysfunction of Microglia during Brain Development: Consequences for Synapses and Neural Circuits. Front Synaptic Neurosci 9:9. doi:10.3389/fnsyn.2017.00009

Paolicelli RC, Sierra A, Stevens B, Tremblay ME, Aguzzi A, Ajami B, Amit I, Audinat E, Bechmann I, Bennett M, Bennett F, Bessis A, Biber K, Bilbo S, Blurton-Jones M, Boddeke E, Brites D, Brône B, Brown GC, Butovsky O, Carson MJ, Castellano B, Colonna M, Cowley SA, Cunningham C, Davalos D, De Jager PL, de Strooper B, Denes A, Eggen BJL, Eyo U, Galea E, Garel S, Ginhoux F, Glass CK, Gokce O, Gomez-Nicola D, González B, Gordon S, Graeber MB, Greenhalgh AD, Gressens P, Greter M, Gutmann DH, Haass C, Heneka MT, Heppner FL, Hong S, Hume DA, Jung S, Kettenmann H, Kipnis J, Koyama R, Lemke G, Lynch M, Majewska A, Malcangio M, Malm T, Mancuso R, Masuda T, Matteoli M, McColl BW, Miron VE, Molofsky AV, Monje M, Mracsko E, Nadjar A, Neher JJ, Neniskyte U, Neumann H, Noda M, Peng B, Peri F, Perry VH, Popovich PG, Pridans C, Priller J, Prinz M, Ragozzino D, Ransohoff RM, Salter MW, Schaefer A, Schafer DP, Schwartz M, Simons M, Smith CJ, Streit WJ, Tay TL, Tsai LH, Verkhratsky A, von Bernhardi R, Wake H, Wittamer V, Wolf SA, Wu LJ, Wyss-Coray T (2022) Microglia states and nomenclature: A field at its crossroads. Neuron 110 (21):3458–3483. doi:10.1016/j.neuron.2022.10.020

Parkhurst CN, Yang G, Ninan I, Savas JN, Yates JR, 3rd, Lafaille JJ, Hempstead BL, Littman DR, Gan WB (2013) Microglia promote learning-dependent synapse formation through brain-derived neurotrophic factor. Cell 155 (7):1596–1609. doi:10.1016/j.cell.2013.11.030

Pawelec P, Ziemka-Nalecz M, Sypecka J, Zalewska T (2020) The Impact of the CX3CL1/CX3CR1 Axis in Neurological Disorders. Cells 9 (10). doi:10.3390/cells9102277

Paxinos G, Watson C (2007) The rat brain in stereotaxic coordinates. 6th edn. Academic Press/Elsevier, Amsterdam ; Boston ;

Peng Y, Guo G, Shu B, Liu D, Su P, Zhang X, Gao F (2017) Spinal CX3CL1/CX3CR1 May Not Directly Participate in the Development of Morphine Tolerance in Rats. Neurochem Res 42 (11):3254–3267. doi:10.1007/s11064-017-2364-z

Robinson SP, Langan-Fahey SM, Johnson DA, Jordan VC (1991) Metabolites, pharmacodynamics, and pharmacokinetics of tamoxifen in rats and mice compared to the breast cancer patient. Drug Metab Dispos 19 (1):36–43

Rochefort H, Borgna JL (1981) Differences between oestrogen receptor activation by oestrogen and antioestrogen. Nature 292 (5820):257–259. doi:10.1038/292257a0

Sadek SM, Khatri SN, Kipp Z, Dunn KE, Beckmann JS, Stoops WW, Hinds TD, Jr., Gipson CD (2024) Impacts of xylazine on fentanyl demand, body weight, and acute withdrawal in rats: A comparison to lofexidine. Neuropharmacology 245:109816. doi:10.1016/j.neuropharm.2023.109816

Scofield MD, Heinsbroek JA, Gipson CD, Kupchik YM, Spencer S, Smith AC, Roberts-Wolfe D, Kalivas PW (2016) The Nucleus Accumbens: Mechanisms of Addiction across Drug Classes Reflect the Importance of Glutamate Homeostasis. Pharmacol Rev 68 (3):816–871. doi:10.1124/pr.116.012484

Sher ME, Bank S, Greenberg R, Sardinha TC, Weissman S, Bailey B, Gilliland R, Wexner SD (1999) The influence of cigarette smoking on cytokine levels in patients with inflammatory bowel disease. Inflamm Bowel Dis 5 (2):73–78. doi:10.1097/00054725-199905000-00001

Shi FD, Piao WH, Kuo YP, Campagnolo DI, Vollmer TL, Lukas RJ (2009) Nicotinic attenuation of central nervous system inflammation and autoimmunity. J Immunol 182 (3):1730–1739. doi:10.4049/jimmunol.182.3.1730

Soares AR, Picciotto MR (2024) Nicotinic regulation of microglia: potential contributions to addiction. J Neural Transm (Vienna) 131 (5):425–435. doi:10.1007/s00702-023-02703-94

Souter, E. A., Chen, Y. C., Zell, V., Lallai, V., Steinkellner, T., Conrad, W. S., Wisden, W., Harris, K. D., Fowler, C. D., & Hnasko, T. S. (2022). Disruption of VGLUT1 in Cholinergic Medial Habenula Projections Increases Nicotine Self-Administration. eNeuro, 9(1), ENEURO.0481-21.2021. 10.1523/ENEURO.0481-21.2021

Tarnowski BI, Spinale FG, Nicholson JH (1991) DAPI as a useful stain for nuclear quantitation. Biotech Histochem 66 (6):297–302

Wakabayashi-Ito N, Nagata S (1994) Characterization of the regulatory elements in the promoter of the human elongation factor-1 alpha gene. J Biol Chem 269 (47):29831–29837

Wickel J, Chung HY, Ceanga M, von Stackelberg N, Hahn N, Candemir Ö, Baade-Büttner C, Mein N, Tomasini P, Woldeyesus DM, Andreas N, Baumgarten P, Koch P, Groth M, Wang ZQ, Geis C (2024) Repopulated microglia after pharmacological depletion decrease dendritic spine density in adult mouse brain. Glia 72 (8):1484–1500. doi:10.1002/glia.24541

Wieghofer P, Prinz M (2016) Genetic manipulation of microglia during brain development and disease. Biochim Biophys Acta 1862 (3):299–309. doi:10.1016/j.bbadis.2015.09.019

Yi MH, Liu YU, Liu K, Chen T, Bosco DB, Zheng J, Xie M, Zhou L, Qu W, Wu LJ (2021) Chemogenetic manipulation of microglia inhibits neuroinflammation and neuropathic pain in mice. Brain Behav Immun 92:78–89. doi:10.1016/j.bbi.2020.11.030

Yona S, Kim KW, Wolf Y, Mildner A, Varol D, Breker M, Strauss-Ayali D, Viukov S, Guilliams M, Misharin A, Hume DA, Perlman H, Malissen B, Zelzer E, Jung S (2013) Fate mapping reveals origins and dynamics of monocytes and tissue macrophages under homeostasis. Immunity 38 (1):79–91. doi:10.1016/j.immuni.2012.12.001

Zhang B, Zou J, Han L, Beeler B, Friedman JL, Griffin E, Piao YS, Rensing NR, Wong M (2018) The specificity and role of microglia in epileptogenesis in mouse models of tuberous sclerosis complex. Epilepsia 59 (9):1796–1806. doi:10.1111/epi.14526

Zhao XF, Alam MM, Liao Y, Huang T, Mathur R, Zhu X, Huang Y (2019) Targeting Microglia Using Cx3cr1-Cre Lines: Revisiting the Specificity. eNeuro 6 (4). doi:10.1523/ENEURO.0114-19.2019

